# Comprehensive top-down mass spectral repository enables pan-dataset analysis and top-down spectral prediction

**DOI:** 10.64898/2026.02.20.707032

**Authors:** Kun Li, Kaiyuan Liu, James M. Fulcher, Philipp T. Kaulich, Haixu Tang, Xiaowen Liu

## Abstract

Mass spectral libraries have become essential resources for training deep learning (DL) models for spectral prediction and *de novo* sequencing in bottom-up mass spectrometry (BU-MS). Compared with BU-MS, top-down MS (TD-MS) offers unique advantages for characterizing intact proteoforms by analyzing proteoforms without enzymatic digestion. Despite these advantages, large-scale spectral libraries for TD-MS are currently lacking. Here we present TopRepo, the first comprehensive and publicly available repository of TD-MS spectra, comprising more than 12 million spectra acquired from 12 species across seven types of mass spectrometers. Using TopRepo, we constructed a large-scale top-down spectral library containing over 5 million spectra with proteoform and fragment-ion annotations. We demonstrate that TopRepo enables pan-dataset analyses of N-terminal processing, mass shifts, and other proteoform characteristics identified by TD-MS. Furthermore, we show that the TopRepo spectral library substantially improves proteoform identification through spectral library searching and supports the training of DL models for high-accuracy top-down spectral prediction.

## Introduction

Mass spectral libraries are widely used in bottom-up mass spectrometry (BU-MS) for peptide identification through spectral library searching [1], particularly in data-independent acquisition mass spectrometry (DIA-MS) workflows [2]. Numerous large-scale experimental BU-MS spectral libraries have been established, including ProteomeTools [3], NIST [4], and MassIVE-KB [5]. More recently, these spectral resources have become essential for training deep learning (DL) models for bottom-up spectral prediction [6] and *de novo* peptide sequencing [7].

Compared with bottom-up mass spectrometry (BU-MS), top-down mass spectrometry (TD-MS) provides distinct advantages for studying complex proteoforms and combinatorial post-translational modifications (PTMs), as it directly analyzes intact proteins and offers a global view of proteoforms [8]. As in BU-MS, comprehensive top-down spectral libraries are essential for advancing computational methods and DL models for TD-MS data analysis. Access to sufficiently large and diverse top-down spectral libraries enables DL models to learn improved spectral similarity functions that more effectively distinguish true from false spectral identifications [9], and to generate *in silico* top-down spectral libraries that enhance spectral identification [6]. Moreover, well-annotated top-down spectral libraries enable more precise characterization and localization of PTMs. While existing database search tools efficiently map spectra to protein sequences, they frequently exhibit limited confidence in PTM identification and site localization [10]. In contrast, spectral libraries enriched with accurate proteoform annotations provide a robust framework for both comprehensive proteoform characterization and reliable PTM assignment in TD-MS. Despite their utility, large-scale and comprehensive top-down spectral libraries remain largely unavailable.

Here, we present TopRepo, a repository of top-down MS data comprising more than 12 million tandem mass spectrometry (MS/MS) spectra and a spectral library of over 5.1 million spectra with proteoform identifications and fragment ion annotations. To our knowledge, this represents the largest top-down spectral library to date, spanning 12 species and 7 types of mass spectrometers. Leveraging TopRepo, we systematically evaluated spectral identification rates and analyzed proteoform features, including sequence length, N-terminal processing, mass shifts, signal peptides, and reproducibility across datasets. We further demonstrated that DL models trained on the TopRepo spectral library achieved high accuracy in top-down spectral prediction. Finally, we showed that a large-scale library constructed from TopRepo increased spectral library based proteoform identification by 41.5% compared with a library derived from a single MS dataset.

## Results

TopRepo comprises a comprehensive top-down spectral repository constructed from 3,165 MS files associated with 33 publications (**Fig. 1a** and **Table S1, S2**), building upon a previously reported collection of TD-MS datasets [11]. Because five publications included data from two species, whereas the remaining 28 publications each involved a single species, we divided each of the five multi-species studies into two projects and assigned each of the 28 single-species studies to one project, resulting in a total of 38 projects (Supplemental **Table S3**) collected from 87 TD-MS repository datasets (Supplemental **Table S4**). To ensure consistency, each dataset was further partitioned into subdatasets generated using the same sample type, instrument, and fragmentation method, yielding 112 subdatasets (Supplemental **Table S5**).

**Fig. 1:**
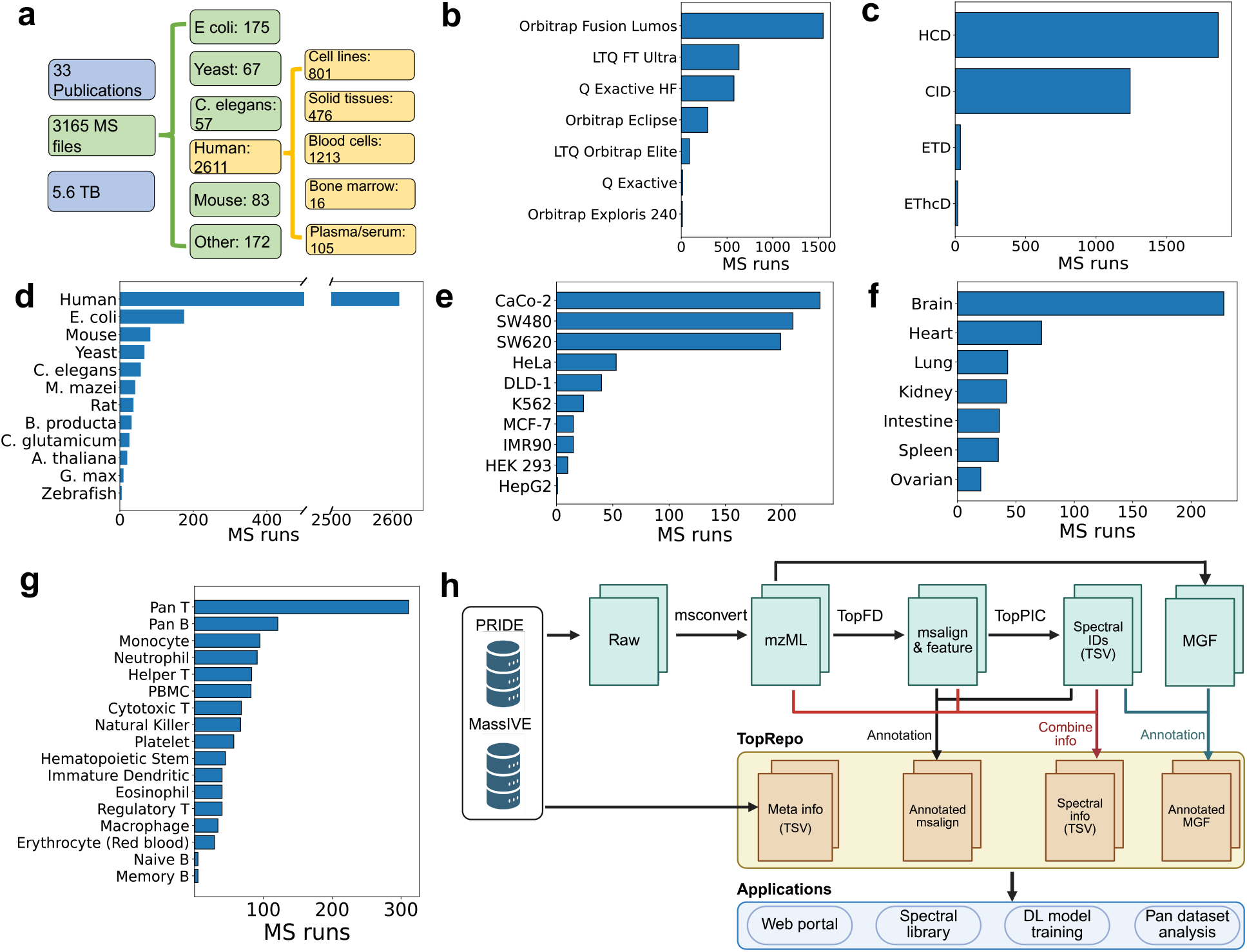
Overview of TopRepo. (a) A total of 3,165 top-down raw MS files were collected to construct TopRepo. Distributions of (b) mass spectrometer platforms and (c) fragmentation methods used to generate these datasets. Summary of MS files derived from (d) 12 species, (e) 10 human cell lines, (f) 7 types of human solid tissues, and (g) 17 types of human blood cells. (h) Data processing workflow for building TopRepo. Raw MS files were downloaded from the PRIDE, MassIVE, and iProX repositories and converted to centroided mzML and MGF formats. Feature detection and spectral deconvolution of mzML files were performed using TopFD, followed by database search-based proteoform identification using TopPIC. Identified proteoforms were used to annotate deconvoluted fragment masses in msalign files and fragment peaks in MGF files. TSV files with comprehensive spectral information were generated by integrating spectral information from mzML and msalign files with proteoform identifications reported by TopPIC. Metadata describing experimental methods were also retrieved from PRIDE, MassIVE, and iProX. TopRepo provides metadata files, annotated msalign files, annotated MGF files, and TSV files containing integrated spectral information. MS data in TopRepo were then used for web portal development, spectral library construction, DL model training, and pan-dataset analyses. Panel (h) was created with BioRender.com.

These data were generated from 12 species using 7 types of mass spectrometers and 4 dissociation methods (**Fig. 1b–d**). Of the 3,165 MS raw files, 2,611 (82.5%) were derived from human samples, encompassing 10 human cell lines (**Fig. 1e**), 7 types of human solid tissues (**Fig. 1f**), 17 types of human blood cells (**Fig. 1g**), 4 types of bone marrow cells, and plasma/serum samples (Supplemental **Fig. S1**).

All spectra were processed using a unified analysis pipeline consisting of msconvert from ProteoWizard [12], TopFD [13], and TopPIC [9], with false discovery rate (FDR)-based quality control to ensure data quality and harmonization (**Fig. 1h** and **Methods**). For raw files acquired with High-Field Asymmetric Waveform Ion Mobility Spectrometry (FAIMS), TopFD generated multiple deconvoluted MS (msalign) files per raw file, resulting in 4,615 msalign files in total (Supplemental **Tables S6, S7**). Altogether, TopRepo contains 12,879,563 MS/MS spectra (Supplemental **Fig. S2**).

### Spectral and proteoform identifications

Of the spectra in TopRepo, we successfully identified 5,147,250 proteoform-spectrum matches (PrSMs) (**Fig. 2a**). We further constructed nonredundant representative proteoform sets at the msalign file level, project level, and across the entire repository, resulting in 1,962,020 proteoform identifications after file-level duplication removal (Supplemental **Table S7**), 329,019 proteoform identifications after project-level duplication removal (Supplemental **Table S3**), and 284,527 unique proteoforms after repository-level duplication removal (**Fig. 2b**). These proteoforms were derived from 17,243 proteins, with the highest counts originating from human, mouse, and *E. coli* (**Fig. 2c**).

**Fig. 2:**
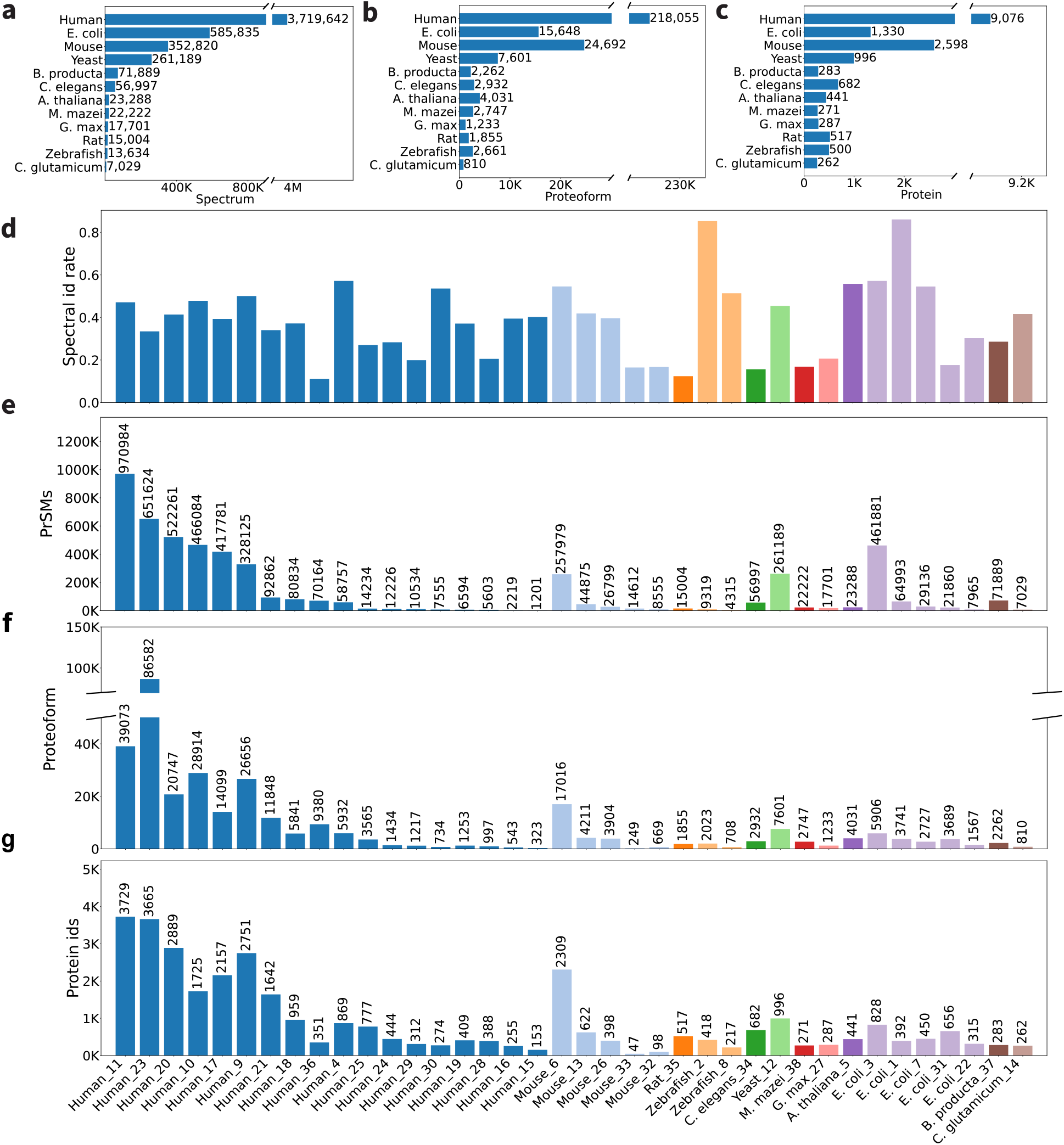
Spectral, proteoform, and protein identifications in TopRepo. (a) Number of spectral (PrSM) identifications, (b) proteoform identifications, and (c) protein identifications across 12 species. (d) Spectral identification rates and numbers of (e) spectral, (f) proteoform, and (g) protein identifications across 38 projects.

We next examined the spectral identification rate, defined as the percentage of acquired spectra with PrSM identifications, as well as the number of spectral (PrSM) identifications for each project (**Fig. 2d**,**e**). The identification rate of all spectra in TopRepo was 40.0%, with substantial variance across projects, highlighting the heterogeneity of the datasets. Project 1 (*E. coli*) and Project 2 (zebrafish) achieved identification rates of 86.1% and 85.3%, respectively, whereas the highest identification rate among more complex human samples was 57.2% (Project 4) (Supplemental **Table S3**). At the proteoform level, the best-performing project (Project 23, the human blood proteoform atlas project [14]) identified 86,582 proteoforms (**Fig. 2f**). At the protein level, Project 11 identified 3,729 proteins, demonstrating the potential of TDP to achieve high proteome coverage in large-scale proteomic studies (**Fig. 2g**).

We also analyzed the length distribution of identified proteoforms across projects (**Fig. 3a**). The average proteoform length was 70.4 amino acids across the 284,527 proteoforms, whereas Projects 25 and 31 reported the longest average lengths (133.4 and 121.3 amino acids, respectively) (Supplemental **Table S3**), both generated using 21 Tesla FT-ICR mass spectrometers. We further categorized proteoform identifications into four classes: complete (no truncations), N-terminal (C-terminal truncation only, N-terminal methionine excision (NME) is allowed), C-terminal (N-terminal truncation only), and internal (both N- and C-terminal truncations) proteoforms (**Fig. 3b**). Only 12.6% of the 284,527 identified proteoforms were complete. Projects 19 and 30 reported the highest proportions of complete proteoforms (≥59%), although each identified fewer than 1,300 proteoforms. Notably, the average proteoform length was positively correlated with the proportion of complete proteoforms (**Fig. 3c**). Many observed truncation sites of truncated proteoforms matched the pattern of trypsin digestion (**Fig. 3d**,**e** and Supplemental **Fig. S3**,**S4**), consistent with previous findings [11].

**Fig. 3:**
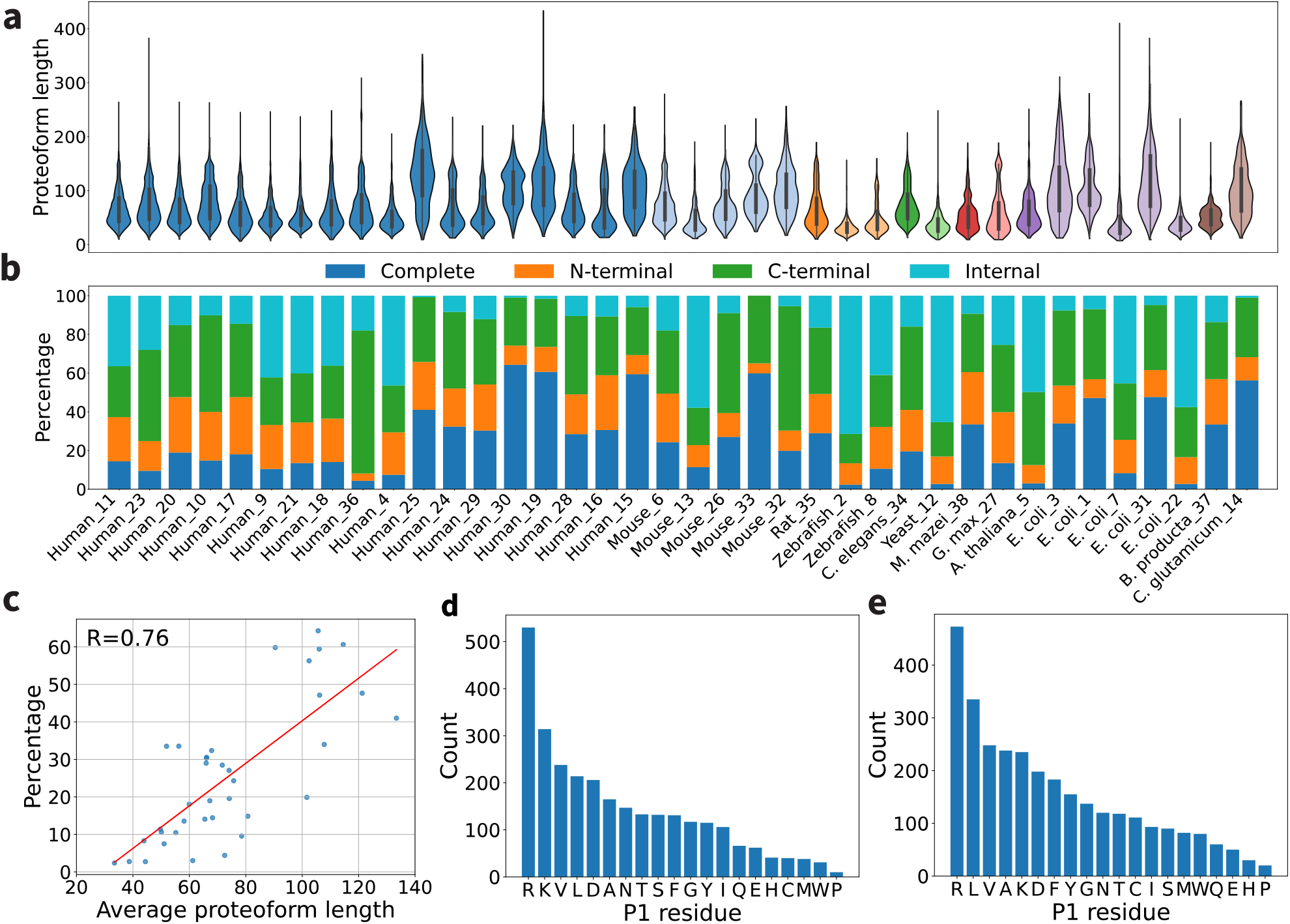
Proteoform lengths and truncation patterns across 38 projects. (a) Distribution of proteoform sequence lengths. (b) Proportions of complete, N-terminal, C-terminal, and internal proteoforms. (c) Pearson correlation between the average proteoform length and the proportion of complete proteoforms across projects. (d) Distribution of P1 amino acid residues (the residue on the N-terminal side of the truncation site) at the C-termini of N-terminal and internal human proteoforms without mass shifts. (e) Distribution of P1 amino acid residues at the N-termini of C-terminal and internal human proteoforms without mass shifts.

### N-terminal methionine excision and N-terminal acetylation

We analyzed the N-terminal forms of 3,481 identified human proteins using complete or N-terminal proteoforms without mass shifts. We restricted the analysis to proteoforms without unknown mass shifts, as mass shifts may introduce errors in the N-terminal forms reported by TopPIC [9]. Specifically, we examined how the P1′ residue (the second amino acid of the protein) influences the rates of NME and N-terminal acetylation (NTA), which are regulated by N-terminal acetyltransferases (NATs). We first compared proteins containing NME proteoforms with those containing non-NME proteoforms. Proteins with P1′ residues A, S, P, T, G, V, or C were predominantly associated with NME proteoforms, whereas proteins with other P1′ residues were primarily associated with non-NME proteoforms (**Fig. 4a**), and some proteins have both NME and non-NME proteoforms. These observations are consistent with the substrate specificities of methionine aminopeptidase 1 (MAP1; A, S, C, G, P) and MAP2 (T and V) [15], as well as previous findings from bottom-up MS studies [16].

**Fig. 4:**
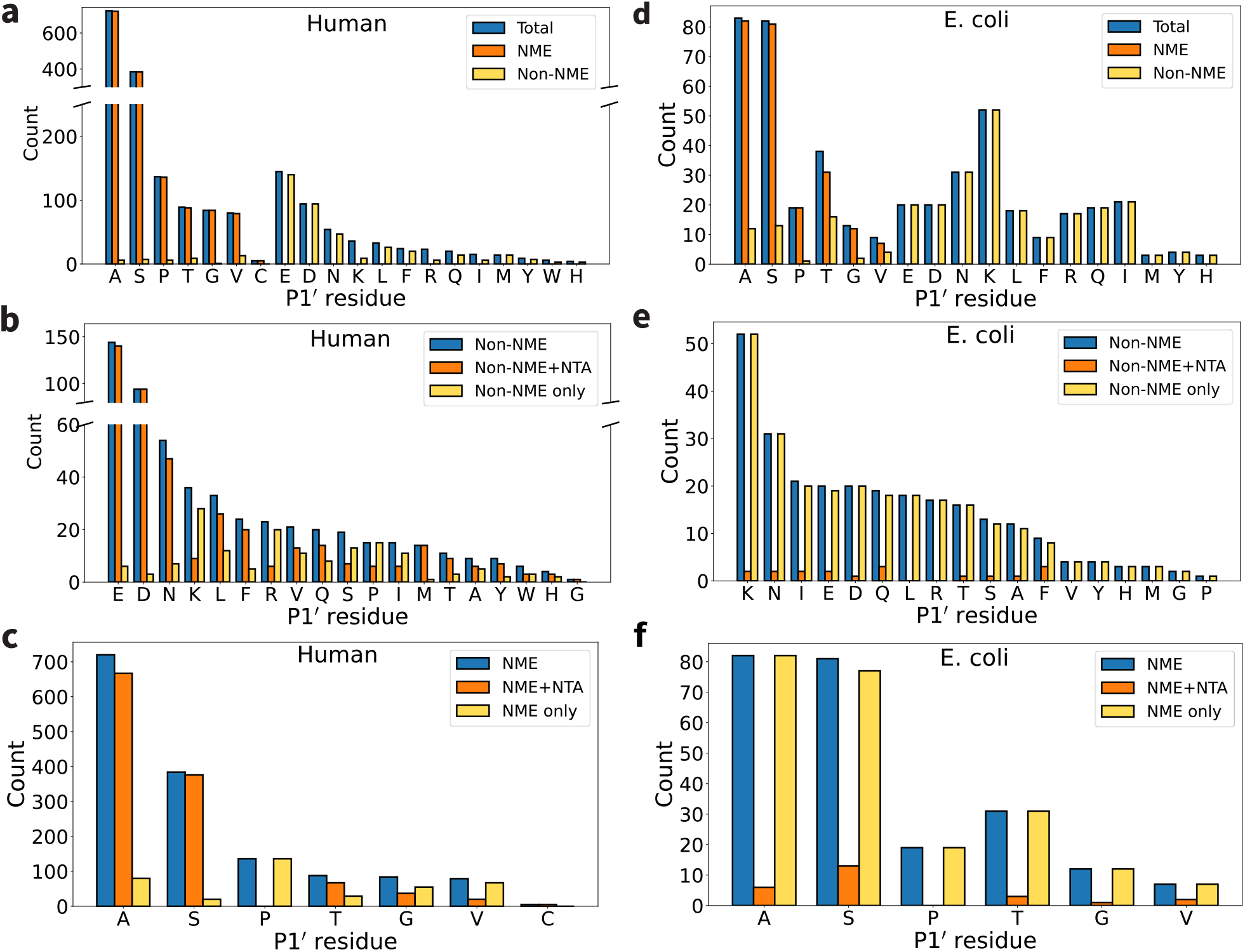
N-terminal processing of human and *E. coli* complete and N-terminal proteoforms without unexpected mass shifts in TopRepo. Comparison of all proteins with a proteoform, proteins with a proteoform with N-terminal methionine excision (NME), and proteins with a proteoform without NME for human (a) and *E. coli* (d). Comparison of all proteins with a proteoform with no NME, proteins with a proteoform with no NME and with NTA, proteins with a proteoform with no NME and no NTA for human (b) and *E. coli* (e). Comparison of all proteins with a proteoform with NME, proteins with a proteoform with NME and NTA, proteins with a proteoform with NME and no NTA for human (c) and *E. coli* (f).

We examined proteins with non-NME proteoforms and compared those with and without NTA (**Fig. 4b)**. Proteins with P1′ residues E, D, N, F, M, T, or Y were predominantly acetylated, consistent with modification by NATs [17, 18], whereas proteins with P1′ residues K or R were largely non-acetylated. We further compared proteins containing proteoforms with both NME and NTA to those with NME only. And proteins with P1′ residues A and S were predominantly acetylated following NME, whereas proteins with P1′ residue P were largely non-acetylated (**Fig. 4c**) [17, 18].

We also investigated the N-terminal forms of *E. coli* proteins using complete and N-terminal proteoforms without mass shifts. The NME patterns in *E. coli* were similar to those observed in human proteins (**Fig. 4d–f**). However, the frequency of NTA in *E. coli* was substantially lower than in human proteins.

### Signal peptides

Protein annotations for 20,405 entries in the UniProt human proteome database (version January 7, 2026) [19] were downloaded, among which 3,603 proteins were annotated with signal peptides. Comparison of these annotations with human proteoform identifications in TopRepo revealed that 207 proteins contained proteoforms whose N-terminal truncations matched the annotated signal peptide cleavage sites (Supplemental **Table S8**). These results demonstrate that TDP can effectively identify signal peptide processing events in proteins.

### Proteoform mass shifts and PTMs

Of the 284,527 proteoforms in TopRepo, 26.0% were unmodified, 5.5% contained only NTA, 56.3% contained only mass shifts, and 12.2% contained both NTA and mass shifts (**Fig. 5a** and Supplemental **Fig. S5**). Among human proteoforms, the most frequently observed mass shifts were matched to oxidation, acetylation, dimethylation, phosphorylation, and metal adducts (Na, K, and Fe[III]) (**Fig. 5c**), consistent with commonly reported PTMs in bottom-up MS studies [10]. These PTMs were also frequently detected in other species (Supplemental **Fig. S6**). Some uncommon mass shifts observed in reported proteoforms may be incorrect. Precursor deconvolution and assignment errors are common in TD-MS data analysis and represent a major source of incorrect mass shifts in identified proteoforms [20].

**Fig. 5:**
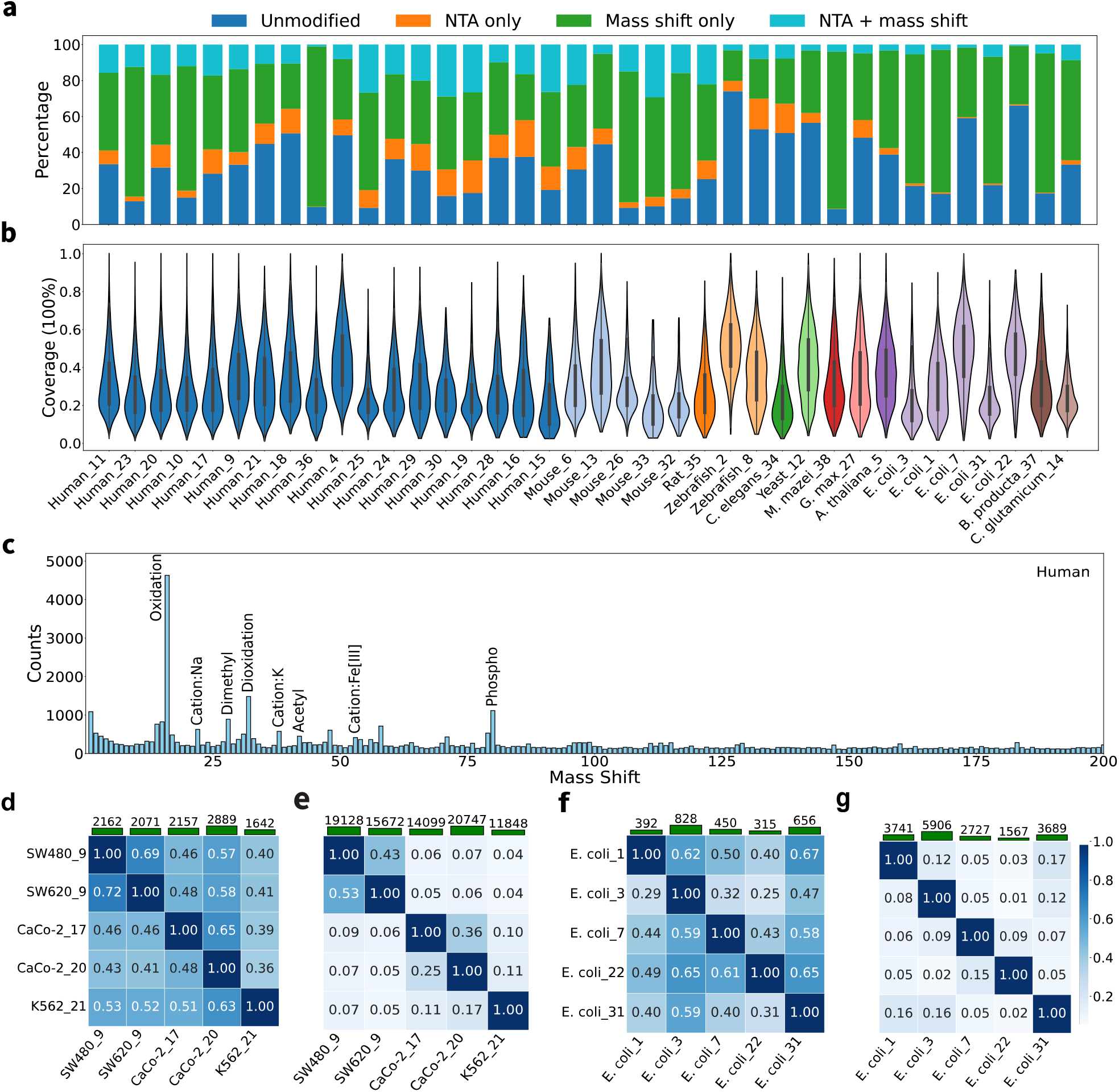
Mass shifts, sequence coverage, and reproducibility of proteoform identifications in TopRepo. (a) Percentages of proteoform identifications without modifications, with only NTA, with only mass shifts, and with both NTA and mass shifts across 38 projects. (b) Distribution of proteoform sequence coverage across the 38 projects. (c) Distribution of observed mass shifts in human proteoform identifications within the range of 0–200 Da with a bin size of 1 Da. Heat maps showing overlaps of (d) protein and (e) proteoform identifications across 5 TD-MS datasets from 4 human cell lines. Heat maps showing overlaps of (f) protein and (g) proteoform identifications across 5 TD-MS datasets from *E. coli* samples. The numbers on the green bars indicate the total identifications for each dataset. Each value in the heat map represents the ratio of shared identifications between two datasets to the total number of identifications in the dataset corresponding to the row.

We further examined proteoform sequence coverage, which is critical for confident PTM localization. The average sequence coverage across the 284,527 proteoform identifications was approximately 30.9% (**Fig. 5b** and Supplemental **Fig. S7**). Of the 284,527 proteoform identifications, 89,731 were unmodified or contained only NTA, with an average sequence coverage of 37.6%, whereas the remaining 194,796 proteoform identifications contained unknown mass shifts and exhibited a lower average sequence coverage of 27.9%, highlighting current limitations of TDP for precise PTM localization. Notably, besides the dissociation technique, sequence coverage is also influenced by the performance of spectral deconvolution algorithms, so improvements in deconvolution methods are expected to enhance coverage and PTM localization accuracy.

### Proteoform identification by different datasets

We assessed the reproducibility of proteoform and protein identifications across multiple human cell line datasets. Protein identifications from five TDP datasets representing four human cell lines, each with at least 1,000 identified proteins, were first compared (**Fig. 5e**). All the datasets were generated using Orbitrap mass spectrometers. Pairwise comparisons showed that 36%–72% of protein identifications were shared between datasets. Because TDP preferentially identifies high-abundance proteins, independent TDP experiments of human cell lines tend to identify a similar set of abundant proteins. The highest protein-level reproducibility (72%) was observed between the SW480 and SW620 datasets generated within the same project (Project 9).

In contrast, proteoform-level comparisons revealed substantially larger variability across experiments conducted by different laboratories (**Fig. 5f**). The highest proteoform-level overlap (53%) occurred between the SW480 and SW620 datasets. The two CaCo-2 datasets generated by different projects also showed moderate reproducibility (36%). However, pairwise overlaps among other datasets were ≤17%, indicating limited cross-study consistency at the proteoform level.

To investigate the reasons for low reproducibility, we compared proteoform identifications of protein PHB1 in CaCo-2 cells from Project 17 and Project 20 (18 proteoforms identified in each dataset; Supplemental **Table S9**). Only 7 proteoforms were shared between the two projects. Similarly, comparison of HINT1 proteoforms identified in SW480 cells (Project 9; 21 proteoforms) and CaCo-2 cells (Project 17; 16 proteoforms) showed that only 2 proteoform was shared (Supplemental **Table S10**). The primary reason for low reproducibility was the identification of numerous truncated proteoforms unique to individual datasets. Consistently, Kaulich et al. [21] reported that truncated proteoform identifications differ markedly among TD-MS sample preparation protocols. A similar pattern was observed in five TDP datasets of *E. coli* samples (**Fig. 5g**,**h**), where pairwise proteoform-level reproducibility was ≤17%.

### Comparison of protein abundances measured by TDP and BUP

We compared protein abundances reported by TDP and BUP across four human cell lines: HeLa, HEK 293, MCF-7, and K562. Protein abundances for BUP were obtained from a previous study [22], where abundances were estimated using iBAQ values averaged across three replicates for each dataset. For TDP, protein abundance was estimated as the sum of signal intensities of all identified proteoforms corresponding to each protein (**Methods**). It should be noted that TDP-derived abundances are not absolute, as they are influenced by proteoform ionization efficiencies. Overall, TDP and BUP showed consistent protein abundance trends, with Pearson correlation coefficients (PCCs) ranging from 0.40 to 0.41 (Supplemental **Fig. S8**). The moderate correlation likely reflects differences in proteoform/peptide separation, variability in ionization efficiencies, and other experimental factors.

### Improved spectral library-based proteoform identification

Our previous study demonstrated that limited spectral library size is a major constraint on achieving high sensitivity in top-down spectral library searches [6]. We therefore evaluated whether the large-scale library derived from TopRepo could improve spectral identification performance. We constructed two human higher-energy collisional dissociation (HCD) spectral libraries of different sizes. The first library, referred to as SW480-2D, was generated from the first replicate of the SW480 top-down MS dataset with two-dimensional (2D) separation in Project 9. The second library, referred to as HUMAN-HCD, was constructed using all human HCD spectra in TopRepo, excluding the second replicate of the SW480 2D dataset from Project 9 (**Methods**). The SW480-2D library contained 5,360 spectra corresponding to 3,876 proteoforms, whereas the HUMAN-HCD library comprised 253,615 spectra representing 136,758 proteoforms.

We then searched the second replicate of the SW480 2D dataset (22,924 MS/MS spectra) against each library separately using TopLib [6] (**Methods**). Compared with the SW480-2D library, the HUMAN-HCD library increased spectral identifications by 19.3% (10,329 vs. 12,325) at a cosine similarity cutoff of 0.3 and by 17.0% (14,262 vs. 16,690) at a 1% spectrum-level FDR. At the proteoform level, the HUMAN-HCD library increased identifications by 41.5% compared with the SW480-2D library (3,148 vs 4,455) with the cosine similarity cutoff of 0.3. Among the 4,455 proteoforms, 3,739 were also supported by database search, whereas 716 were uniquely identified through spectral library searching (Supplemental **Fig. S9**). These results demonstrate that the large-scale spectral library derived from TopRepo substantially enhances proteoform identification sensitivity in top-down spectral library searches.

### Top-down spectral prediction

Building upon PredFull [23], previously developed for peptide MS/MS spectral prediction, we developed **TD-Pred** (**Fig. 6a**,**b** and **Methods**), a DL model that integrates convolutional neural networks (CNNs) and transformer layers for top-down spectral prediction using proteoform sequences without mass shifts or PTMs. Proteoform sequences are encoded using one-hot encoding combined with residue mass, positional, and length features. This encoding is processed by a CNN subnetwork consisting of eight parallel modules with kernel sizes ranging from 2 to 9, enabling the model to capture local sequence dependencies up to four residues on each side. The CNN outputs are concatenated with the original sequence encoding. In addition, meta-information is encoded and appended to each column of the sequence matrix (**Fig. 6a**), allowing the transformer layers to access global meta-features at every sequence position. The final representation is input into a transformer architecture comprising six encoder and six non-autoregressive decoder layers for spectral prediction (**Fig. 6b**).

**Fig. 6:**
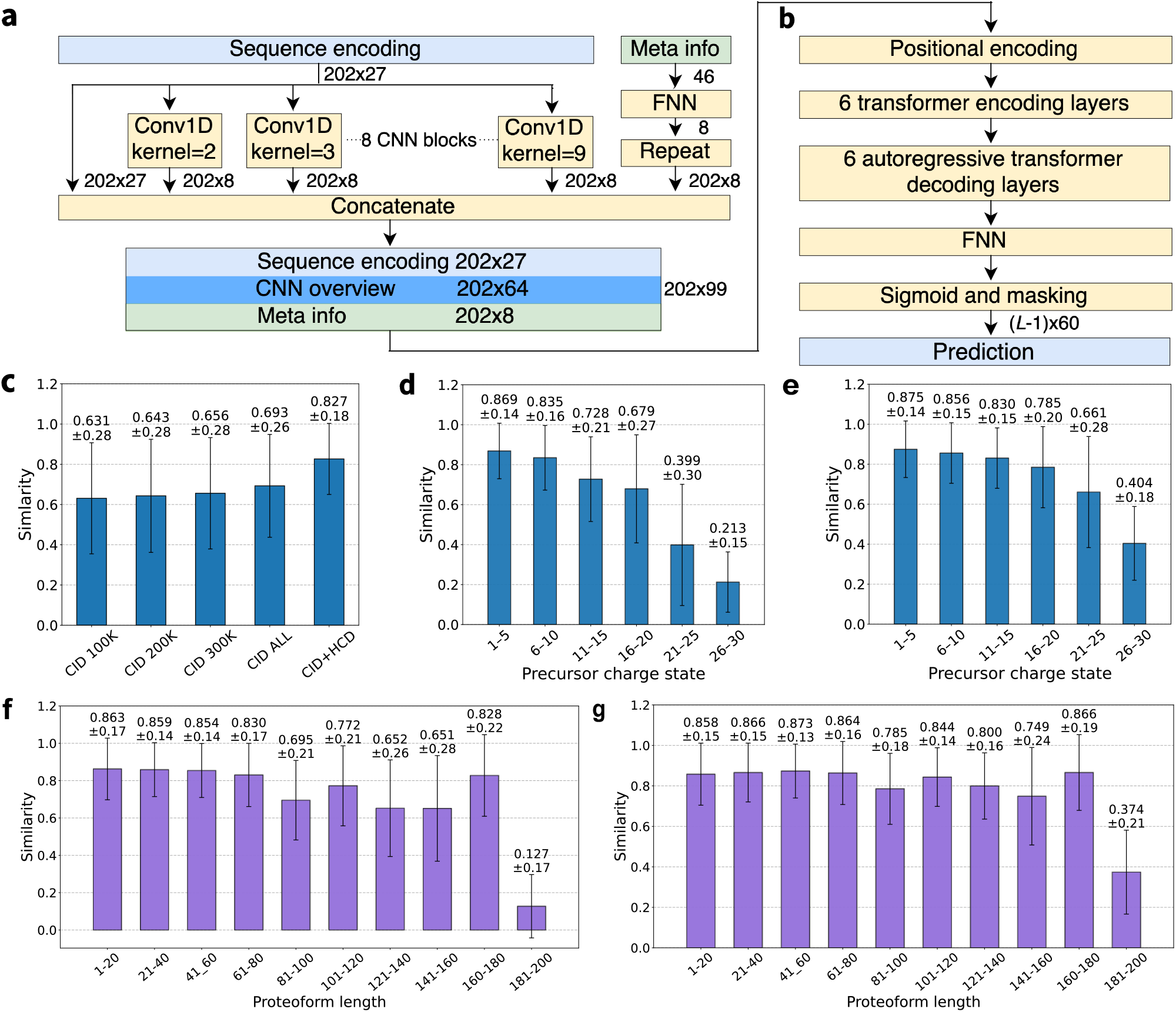
Overview and evaluation of the TD-Pred model. (a) Sequence encoding scheme. Each amino acid in the padded sequence (length 202) is represented by a 27-dimensional vector comprising a 24-dimensional one-hot encoding and three numerical features: residue mass, normalized position, and normalized sequence length. To capture local sequence context, a subnetwork of eight convolutional blocks (each containing a single convolutional layer with kernel sizes from 2 to 9) is applied, and the resulting local features are appended to the sequence encoding. An embedded meta-information vector is added to each residue position, enabling the spectral prediction network to access experimental metadata throughout the sequence. (b) Transformer-based spectral prediction architecture. The encoded sequence is processed by a positional encoding layer, six encoder layers, and six non-autoregressive decoder layers, followed by a feed-forward network and masking layer. The model outputs a backbone representation of the spectrum as an (*L* − 1) × 60 matrix, where *L* is the input sequence length. (c) Prediction accuracy on the VAL-CID dataset using 5 training datasets of different sizes with the backbone representation. Prediction accuracy on the VAL-ALL dataset across proteoforms with varying precursor charge states (d) and sequence lengths (f) using the backbone representation. Prediction accuracy on the VAL-ALL dataset across proteoforms with varying precursor charge states (e) and sequence lengths (g) using the simplified representation.

Deconvoluted top-down MS/MS spectra are represented using common N-terminal and C-terminal fragment ions (**Methods**) [23]. For a proteoform of length *L*, the spectrum is encoded as an (*L*–1) × 60 matrix, referred to as the backbone representation, where *L*–1 corresponds to the number of peptide bonds. Each row contains 60 intensity values representing the relative abundances of the common N-terminal and C-terminal fragment ions across charge states 1–30. We also evaluated a simplified representation, in which a spectrum is encoded as an (*L*–1) × 2 matrix, in which the two rows contain the total abundance of N- and C-terminal fragment ions summed across charge states.

We first trained and evaluated TD-Pred using 472,208 top-down collision-induced dissociation (CID) MS/MS spectra acquired on a Thermo Fusion Lumos instrument from TopRepo. To assess spectral reproducibility, we computed cosine similarity between backbone representations of 10,000 spectral pairs generated from the same proteoform and precursor charge state. The average cosine similarity was 0.946 and similarity scores were consistent across proteoforms of varying lengths (Supplemental **Fig. S10**).

Next, we evaluated prediction performance as a function of training set size using the backbone representation. The dataset was split into 376,497 training spectra and 95,711 validation spectra. Increasing the training set from 100,000 to 376,497 spectra improved validation cosine similarity from 0.631 to 0.693 (**Fig. 6c**). To further enhance performance, we combined CID and HCD spectra acquired on Thermo Fusion Lumos and Q Exactive HF instruments (TRAIN_ALL: 601,520 spectra; VAL_ALL: 152,124 spectra). This strategy substantially improved prediction accuracy to 0.827 (**Fig. 6c**) and reduced the standard deviation of cosine similarity from 0.28 to 0.18. An example predicted spectrum with cosine similarity of 0.938 is shown in Supplemental **Fig. S11**.

We further examined prediction accuracy across precursor charge states and proteoform lengths using TRAIN_ALL and VAL_ALL (**Fig. 6d,f**). Prediction accuracy decreased with increasing charge state, from 0.869 for charge states 1 – 5 to 0.213 for charge states 26 – 30 (**Fig. 6d**). This decline likely reflects limited training spectra with high charge states (Supplemental **Fig. S12a**) and the increased complexity of fragment ion distributions at higher charges. Similarly, prediction accuracy decreased for proteoforms longer than 80 residues, with a pronounced drop for lengths of 180 – 200 (**Fig. 6f**). Long proteoforms are underrepresented in the training data (**Supplemental Fig. S12b**), and their spectra are inherently more complex, both contributing to reduced performance. In addition, CID spectra showed slightly higher prediction accuracy than HCD spectra (Supplemental **Fig. S13a**).

Finally, we evaluated the impact of fragment ion charge state prediction on overall accuracy. We modified the prediction objective from the backbone representation to the simplified representation, removing fragment charge states from the prediction task. This modification improved prediction accuracy from 0.836 to 0.865 for CID spectra and from 0.810 to 0.841 for HCD spectra, while also slightly reducing variance in cosine similarity (Supplemental **Fig. S13b**). Notably, the improvement was more significant for spectra from long proteoforms and with high charge states (**Fig. 6f,g**).

## Discussion and Conclusions

We present TopRepo, a large-scale top-down spectral repository encompassing 12,879,563 MS/MS spectra from 12 species, acquired across 7 mass spectrometer platforms and 4 fragmentation methods, together with a spectral library of 5,147,250 annotated spectra. TopRepo is also supported by a dedicated website for open data dissemination. We demonstrate that this large-scale resource substantially improves the training of DL models for *in silico* top-down spectral prediction, and that expanded spectral coverage provided by TopRepo significantly enhances proteoform identification through spectral library searching.

Despite its scale and utility, TopRepo has several limitations that provide avenues for future refinement. First, because deconvoluted spectra may inherit errors introduced during spectral deconvolution, further curation and improved deconvolution algorithms will be essential to enhance data quality. Such curation includes filtering deconvoluted masses using quality metrics, constraining proteoform mass shifts to those consistent with common PTMs and known mutations, and cross validating identifications across multiple spectral deconvolution and identification methods. Second, the repository is currently restricted to spectra acquired on Orbitrap and Fourier-transform ion cyclotron resonance (FT-ICR) platforms. A key future objective is to expand the repository to include TD-MS data generated using time-of-flight (TOF) and other mass spectrometry platforms. Third, since proteoform annotations rely on database search-based identification tools, only approximately 40.0% of spectra are currently annotated. Among the annotated spectra, about 64.6% contain unknown mass shifts, resulting in incomplete or potentially ambiguous proteoform assignments. The development of advanced proteoform separation techniques and MS acquisition methods for higher-quality data, together with improved bioinformatics tools for more accurate proteoform identification, annotation, and PTM localization, will be critical for the continued evolution of TopRepo.

While TD-Pred was optimized for unmodified proteoforms; extending its architecture to accommodate diverse PTMs is a primary future direction. Beyond fragmentation patterns, incorporating additional predictive targets, such as retention or migration time, could significantly enhance DL model utility. Furthermore, because TD-Pred and current spectral library search tools primarily rely on deconvoluted spectra, future investigations leveraging non-deconvoluted spectral data may enable *in silico* prediction of full spectra and improve the confidence of spectral library search based proteoform identifications.

## Methods

### Top-down MS data analysis

All top-down MS/MS datasets were downloaded from ProteomeXChange [24], MassIVE [25] and iProX [26]. A total of 544 raw files in the datasets were excluded from TopRepo (Supplemental **Table S2**) for several reasons, including runs with no samples analyzed, absence of MS/MS spectra, bottom-up MS experiments, MS runs with medium-resolution MS1, selected ion monitoring (SIM) runs, MS imaging data, and errors encountered during data analysis. Metadata describing sample types, mass spectrometer platforms, and fragmentation methods were collected from the corresponding repositories (Supplemental **Table S5**).

Raw MS data files from each experiment were first converted to centroided mzML files using msconvert (version 3.0.10765) in ProteoWizard [12]. The mzML files were subsequently analyzed using TopFD [13] (version 1.7.10), which performed spectral deconvolution, monoisotopic mass assignment of precursor and fragment ions, and proteoform feature detection. The resulting deconvoluted MS/MS spectra were stored in msalign format. Detailed TopFD parameter settings are provided in Supplemental **Table S11**.

Deconvoluted spectra in msalign format were searched against the corresponding UniProt proteome database (see details in Supplemental **Table S12**) concatenated with a decoy database with the same size using TopPIC [9] (version 1.7.11) to identify PrSMs. In TopPIC, precursor and fragment mass tolerances were set to 10 ppm, the maximum number of unexpected modifications (±500 Da) was set to one, NME, NTA, and, NME with NTA were set as variable PTMs, and fixed modifications were specified according to sample preparation protocols. PrSMs were filtered at a 1% spectrum-level Q-value, and identifications were stored in TSV files. PrSMs identified from an msalign file were merged into proteoform groups: two PrSMs were assigned to the same group if they originated from the same proteoform feature or from the same protein with a precursor mass difference <2.2 Da. The PrSM with the best E-value in each group was selected as the representative PrSM. If other PrSMs in the group had inconsistent proteoform identifications, they were replaced with the representative identification. Representative PrSMs were subsequently filtered at a 1% proteoform-level Q-value. Detailed TopPIC parameters are summarized in Supplemental **Table S13**.

Representative PrSMs were further merged at the project and repository levels. Two PrSMs were grouped together if they were derived from the same protein, their overlapping sequence region covered ≥70% of the shorter proteoform, and their precursor mass difference was <2.2 Da. The PrSM with the best E-value in each group was retained as the representative proteoform.

TopRepo stores MS/MS data in four types of files (**Fig. 1h**): (1) TSV files containing experimental metadata; (2) MGF files containing centroided MS/MS peak lists and annotations; (3) msalign file containing deconvoluted precursor and fragment masses with intensities and annotations; and (4) TSV files containing integrated spectral and proteoform identification information extracted from mzML, msalign, and identification outputs. Links for raw MS files are also provided for each dataset.

### Annotation of msalign files

For each deconvoluted MS/MS spectrum, which contains a list of fragment masses, and their intensities and charge states, fragment ion annotation was performed using the following procedure. (1) For each spectrum, theoretical fragment ion masses were computed based on the corresponding proteoform sequence. Fixed modifications and any unexpected mass shifts identified in the proteoform were incorporated into the mass calculations. (2) An experimental deconvoluted fragment mass was considered matched to a theoretical fragment mass if the mass error was within 20 ppm or 0.01 Da. Fragment masses with ±1.00235 Da shifts, which are frequently introduced during spectral deconvolution, were also allowed in the matching with the error tolerance 20 ppm or 0.01 Da. (3) Matched experimental fragments were annotated with the corresponding ion type and cleavage position in the proteoform sequence.

### Annotation of MGF files

Each centroided MS/MS spectrum stored in MGF files contains a list of peaks represented by their mass-to-charge (*m/z*) ratios and intensities. Each centroided spectrum was annotated using its corresponding deconvoluted fragment masses generated by TopFD [13]. When a proteoform identification was obtained by database search using TopPIC [9], the identified proteoform sequence was also used for annotation. The annotation method consisted of the following steps. (1) For each deconvoluted mass, the theoretical isotopic envelope was calculated using the Averagine model [27]. The intensity of each theoretical isotopic peak was determined based on the total abundance of the corresponding deconvoluted mass. (2) The theoretical isotopic peaks were matched to experimental peaks using a mass tolerance of 20 ppm. If an experimental peak matched multiple theoretical peaks, the match with the smallest mass error was retained. (3) Among all experimental peaks matched to the theoretical isotopic envelope, the minimum intensity was selected as a cutoff. Theoretical peaks with intensities lower than this cutoff were discarded. (4) Each remaining matched experimental peak was annotated with the monoisotopic mass and charge state of the corresponding envelope, as well as the theoretical peak mass, *m/z* value, relative intensity, and mass difference with respect to the monoisotopic peak. (5) If a deconvoluted mass was matched to a theoretical fragment mass of the identified proteoform, all experimental isotopic peaks associated with the corresponding theoretical envelope were annotated with the fragment ion type, cleavage position, theoretical fragment mass, and the mass error between experimental and theoretical fragment masses.

### Correlation between BUP and TDP protein abundances

Protein abundances for BU-MS in HeLa, HEK293, MCF-7, and K562 cells were obtained from a previous study [22], which reported log-transformed iBAQ values from three biological replicates for each cell line. Protein abundance was defined as the average of the log-transformed iBAQ values across the three replicates. For TD-MS, a single dataset with the largest number of proteoform identifications was selected for each cell line (Supplemental **Table S14**). Specifically, single-run MS datasets were used for HeLa and HEK293, whereas MS datasets with 2D separation were selected for MCF-7 (8 fractions) and K562 (4 fractions). TD-MS protein abundance was estimated by summing the signal intensities of all identified proteoforms corresponding to each protein across all associated MS files. Only proteins identified by both TD-MS and BU-MS were included in the Pearson correlation coefficient (PCC) analysis (Supplemental **Tables S15–S18**).

### The TD-Pred model

Each protein sequence (maximum length 200 residues) was augmented with start and end tokens and zero-padded to a fixed length of 202, and then represented as a 202 × 27 matrix. Each residue was encoded using a 27-dimensional vector consisting of a 24-dimensional one-hot encoding (for 20 canonical amino acids, selenocysteine, ornithine, the start token, and the end token) and three additional numerical features. The first feature represents the normalized residue mass (residue mass divided by 200, yielding values between 0 and 1; set to 0 for non-amino acid symbols). The second feature encodes the normalized residue position within the proteoform. For a residue at position *x* in a proteoform of length *L*, the normalized position is defined as 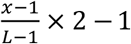, resulting in a values between −1 and 1. The third feature represents the normalized proteoform length (*L*/200).

To capture local sequence patterns, eight CNN modules were applied with kernel sizes ranging from 2 to 9. Each module produced an 8-dimensional representation at each sequence position, resulting in a 64-dimensional local encoding per residue.

A 46-dimensional meta-information vector was constructed and incorporated as the input of the model. This vector included: (1) a 30-dimensional one-hot encoding of the precursor charge state (maximum charge 30), (2) a 9-dimensional one-hot encoding of the instrument type, (3) a 5-dimensional one-hot encoding of the fragmentation type, (4) a normalized proteoform mass (mass/30,000), and (5) a normalized collision energy (NCE) value (NCE/100). The meta-information vector was embedded into an 8-dimensional representation using a fully connected layer, and the resulting embedding was added to each residue position.

After concatenating the CNN-derived 64-dimensional encoding vector with the 27-dimensional residue representation and the 8-dimensional meta-information embedding vector, each residue was represented by a 99-dimensional vector. Therefore, the full input to the model was a 202 × 99 matrix.

This matrix was processed by a transformer encoder–decoder architecture comprising six encoder layers and six non-autoregressive decoder layers. The output of the transformer module was passed through a fully connected layer with a sigmoid activation function to produce the final predictions.

### Model training and validation

Spectra generated from two types of mass spectrometers: Orbitrap Fusion Lumos and Q Exactive HF were used for model training and validation. Spectra of the two types of instruments with proteoform annotations in TopRepo were first filtered using three criteria: (1) the activation method was HCD or CID; (2) the proteoform length was ≤ 200 residues; and (3) the PrSM E-value was ≤ 1 × 10^−4^.

For a proteoform of length *L*, its corresponding experimental spectrum was annotated with b- and y-ions with charge states from 1 to 30. The backbone representation of the spectrum was constructed as a two-dimensional (*L* − 1) × 60 matrix *T*, where the first dimension corresponds to peptide bond cleavage positions and the second dimension represents ion types (b and y ions across 30 charge states each). Each entry *T*[*i, j*] denotes the intensity of the corresponding experimental ion; *T*[*i, j*] = 0 if the ion was not observed. Spectra with fewer than 0.4 × (*L* − 1) nonzero entries in *T* were excluded. A simplified representation was also generated as a (*L* − 1) × 2 matrix, where the two channels correspond to the summed intensities of b and y-ions, respectively, across all charge states at each cleavage position.

The annotated spectra were grouped based on proteoform. Proteoform groups were then split into training and validation sets at an 80:20 ratio, with proteoform level separation to prevent data leakage between the two sets. During splitting, the overall number of spectra was maintained at an approximately 80:20 ratio. The resulting datasets were denoted as TRAIN_ALL and VAL_ALL.

To construct CID-specific datasets, all CID spectra acquired on Orbitrap Fusion Lumos mass spectrometers were extracted from TRAIN_ALL and VAL_ALL, forming TRAIN_CID and VAL_CID. From TRAIN_CID, three smaller training subsets containing 100K, 200K, and 300K spectra were randomly sampled, denoted as TRAIN_CID_100K, TRAIN_CID_200K, and TRAIN_CID_300K, respectively. The spectral prediction model was trained separately using TRAIN_CID_100K, TRAIN_CID_200K, TRAIN_CID_300K, and TRAIN_CID, with VAL_CID used for validation in each case. In addition, TRAIN_ALL and VAL_ALL were used for training and validation, respectively.

The deep learning model TD-Pred was implemented in PyTorch and optimized using the Adam algorithm [28]. Training was performed in two stages. In the first stage, the model was trained for 50 epochs with a batch size of 32 and a learning rate of 3 × 10^−4^. The best-performing model from this stage was further fine-tuned for 10 additional epochs using a reduced learning rate of 5 × 10^−5^. The best model obtained after the second stage was selected as the final model.

### Spectral library building and search

TopLib (version 1.1.0) [6] was used for spectral library construction and library-based identification of top-down MS spectra. To build the HUMAN-HCD spectral library, we collected 1,415,612 MS/MS spectra with proteoform identifications from all human HCD top-down MS runs (1,526 MS files and 1,635 msalign files), excluding the second replicate of the SW480 2D dataset, (six MS files in Project 9), which was used for the evaluation of spectral library search. In total, 258,867 PrSMs were obtained and grouped according to protein accession, precursor mass, charge state, proteoform sequence, instrument type, and activation energy. Two PrSMs were assigned to the same group if (1) they were acquired on the same type of instrument, (2) their NCE values (rounded to integers) were identical, (3) the two proteoforms were from the same protein, (4) the difference in precursor masses was less than 2.2 Da, and (5) their proteoform sequences overlapped by more than 70%. Within each group, the PrSM with the smallest E-value was selected as the representative spectrum for library construction.

For spectral library searching, both precursor and fragment mass tolerances were set to 10 ppm. ±1 Da precursor mass shifts were also permitted to account for common deconvolution errors. Spectral matching was allowed for a spectral pair with different precursor charge states, instrument types, and activation energy values. FDRs were estimated using the target–decoy strategy [6]. Detailed TopLib parameter settings are provided in Supplemental **Table S19**.

### Study funding

This work was supported by the National Institutes of Health (R01CA247863 to X.L. and R01GM130091 to H.T.) and by the National Science Foundation (CCF-2307573 to X.L. and DBI-2011271 to H.T.). A portion of this research was performed by J.M.F. under the National Institutes of Health Common Fund, Human Biomolecular Atlas Program (HuBMAP) grant UG3CA256959, and through the Environmental Molecular Science Laboratory (EMSL, a Department of Energy Office of Science User Facility sponsored by the Office of Biological and Environmental Research program) under Contract No. DE-AC05-76RL01830.

## Acknowledgements

Portions of this research were conducted using high-performance computing resources provided by the Louisiana Optical Network Infrastructure (LONI; http://www.loni.org).

## Author contributions

X.L. and H.T. designed the methods, implemented and reviewed the code, and drafted the manuscript. K. Li collected and processed the data, implemented and trained the TD-Pred model, and contributed to manuscript writing. J.M.F. and P.T.K. collected and processed the data. K. Liu implemented and trained the TD-Pred model. All authors reviewed and approved the final manuscript.

## Code and data availability

The source code for building TopRepo is publicly available at https://github.com/toppic-suite/toprepo. The source code for the TD-Pred model is publicly available at https://github.com/toppic-suite/td-pred. TopRepo data are available at https://www.toprepo.org.

## Notes

The authors used ChatGPT to improve the language and readability of this manuscript. All content was reviewed and edited by the authors, who take full responsibility for the final version of the paper.

## References

1. Cox, J., Prediction of peptide mass spectral libraries with machine learning. Nat Biotechnol, 2023. 41(1): p. 33–43.

2. Lou, R., et al., Benchmarking commonly used software suites and analysis workflows for DIA proteomics and phosphoproteomics. Nat Commun, 2023. 14(1): p. 94.

3. Zolg, D.P., et al., Building ProteomeTools based on a complete synthetic human proteome. Nat Methods, 2017. 14(3): p. 259–262.

4. Yang, X., P. Neta, and S.E. Stein, Extending a Tandem Mass Spectral Library to Include MS(2) Spectra of Fragment Ions Produced In-Source and MS(n) Spectra. J Am Soc Mass Spectrom, 2017. 28(11): p. 2280–2287.

5. Wang, M., et al., Assembling the Community-Scale Discoverable Human Proteome. Cell Syst, 2018. 7(4): p. 412–421 e5.

6. Li, K., H. Tang, and X. Liu, TopLib: Building and Searching Top-Down Mass Spectral Libraries for Proteoform Identification. Anal Chem, 2025. 97(22): p. 11443–11453.

7. Bittremieux, W., et al., Deep Learning Methods for De Novo Peptide Sequencing. Mass Spectrom Rev, 2024.

8. Roberts, D.S., et al., Top-down proteomics. Nat Rev Methods Primers, 2024. 4(1).

9. Kou, Q., L. Xun, and X. Liu, TopPIC: a software tool for top-down mass spectrometry-based proteoform identification and characterization. Bioinformatics, 2016. 32(22): p. 3495–3497.

10. Chen, W., et al., Characterization of Proteoform Post-Translational Modifications by Top-Down and Bottom-Up Mass Spectrometry in Conjunction with Annotations. J Proteome Res, 2023. 22(10): p. 3178–3189.

11. Kaulich, P.T., J.M. Fulcher, and A. Tholey, Properties, Origin, and Consistency of Truncated Proteoforms Across Top-Down Proteomic Studies. Mol Cell Proteomics, 2025. 24(12): p. 101465.

12. Kessner, D., et al., ProteoWizard: open source software for rapid proteomics tools development. Bioinformatics, 2008. 24(21): p. 2534–6.

13. Basharat, A.R., et al., TopFD: A proteoform feature detection tool for top-down proteomics. Anal Chem, 2023. 95(21): p. 8189–8196.

14. Melani, R.D., et al., The Blood Proteoform Atlas: A reference map of proteoforms in human hematopoietic cells. Science, 2022. 375: p. 411.

15. Lowther, W.T. and B.W. Matthews, Structure and function of the methionine aminopeptidases. Biochim Biophys Acta, 2000. 1477(1-2): p. 157–67.

16. Bonissone, S., et al., N-terminal protein processing: a comparative proteogenomic analysis. Mol Cell Proteomics, 2013. 12(1): p. 14–28.

17. Ree, R., S. Varland, and T. Arnesen, Spotlight on protein N-terminal acetylation. Exp Mol Med, 2018. 50(7): p. 1–13.

18. Nguyen, K.T., et al., Control of protein degradation by N-terminal acetylation and the N-end rule pathway. Exp Mol Med, 2018. 50(7): p. 1–8.

19. UniProt, C., UniProt: a worldwide hub of protein knowledge. Nucleic Acids Res, 2019. 47(D1): p. D506–D515.

20. Jeong, K., et al., Precursor deconvolution error estimation: The missing puzzle piece in false discovery rate in top-down proteomics. Proteomics, 2024. 24(3-4): p. e2300068.

21. Kaulich, P.T., et al., Influence of different sample preparation approaches on proteoform identification by top-down proteomics. Nat Methods, 2024. 21(12): p. 2397–2407.

22. Geiger, T., et al., Comparative proteomic analysis of eleven common cell lines reveals ubiquitous but varying expression of most proteins. Mol Cell Proteomics, 2012. 11(3): p. M111 014050.

23. Liu, K., et al., Full-Spectrum Prediction of Peptides Tandem Mass Spectra using Deep Neural Network. Anal Chem, 2020. 92(6): p. 4275–4283.

24. Perez-Riverol, Y., et al., The PRIDE database resources in 2022: a hub for mass spectrometry-based proteomics evidences. Nucleic Acids Res, 2022. 50(D1): p. D543–D552.

25. Choi, M., et al., MassIVE.quant: a community resource of quantitative mass spectrometry-based proteomics datasets. Nat Methods, 2020. 17(10): p. 981–984.

26. Chen, T., et al., iProX in 2021: connecting proteomics data sharing with big data. Nucleic Acids Res, 2022. 50(D1): p. D1522–D1527.

27. Senko, M.W., S.C. Beu, and F.W. McLaffertycor, Determination of monoisotopic masses and ion populations for large biomolecules from resolved isotopic distributions. J. Am. Soc. Mass Spectrom., 1995. 6: p. 229.

28. Kingma, D.P. and J. Ba, Adam: A method for stochastic optimization. arXiv, 2014: p. 1412.6980.

