## Supplemental Figures for "Comprehensive top-down mass spectral repository enables pan-dataset analysis and top-down spectral prediction"

[1]Deming Department of Medicine, School of Medicine, Tulane University, New Orleans, Louisiana, 70112, USA; [2]Luddy School of Informatics, Computing, and Engineering, Indiana University Bloomington, Indiana, 47405, USA; [3]Environmental Molecular Sciences Laboratory, Pacific Northwest National Laboratory, Richland, Washington, 99354, USA; [4]Systematic Proteome Research & Bioanalytics, Institute for Experimental Medicine, Christian-Albrechts-Universität Zu Kiel, Kiel, Germany

### Figures

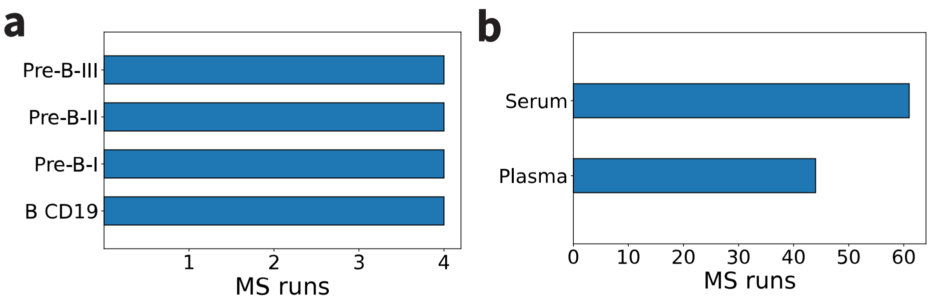


**Fig. S1:** Summary of human top-down MS datasets for bone marrow, plasma, and serum samples in TopRepo. (a) MS files from bone marrow cells. (b) MS files from plasma and serum samples.

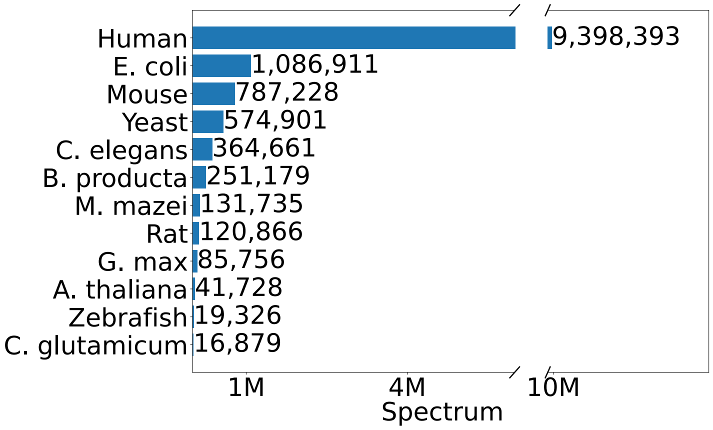


**Fig. S2:** A collection of 12,879,563 top-down MS/MS spectra from 12 species in TopRepo.

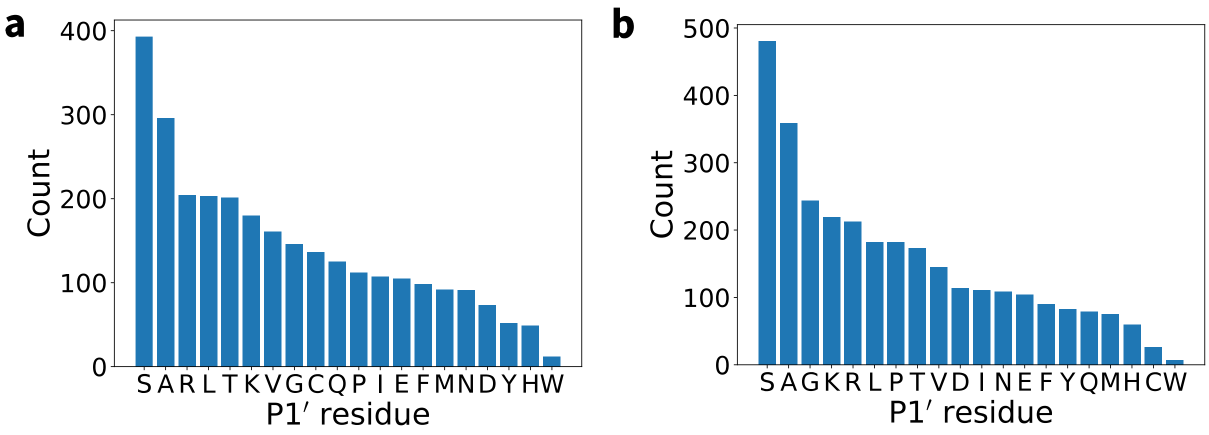


**Fig. S3:** Distributions of the P1' amino acid residues (the residue on the C-terminal side of the truncation site) of truncated human proteoforms without mass shifts. (a) Distributions of the P1' amino acid residues of the C-termini of N-terminal and internal human proteoforms. (b) Distributions of the P1' amino acid residues of the N-termini of C-terminal and internal human proteoforms.

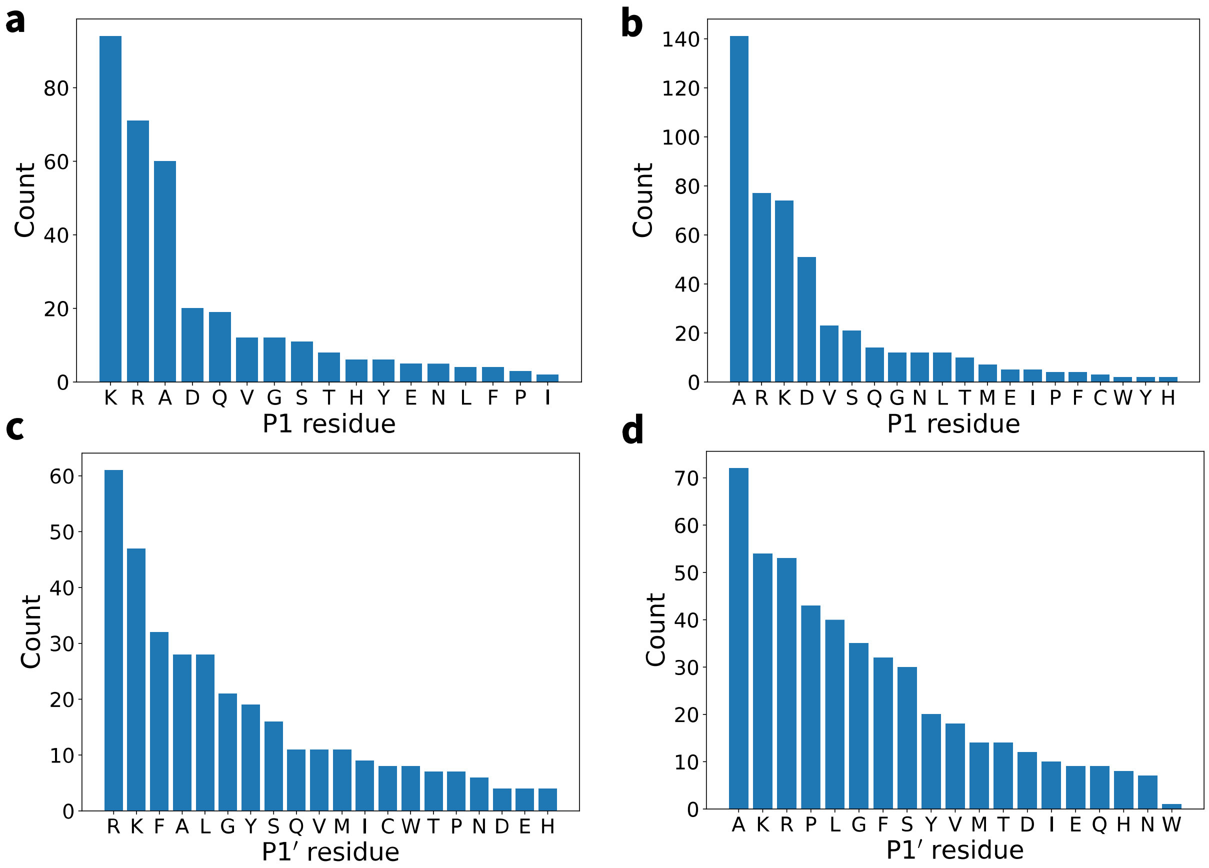


**Fig. S4:** Distributions of the P1 and P1' amino acid residues of truncated *E. coli* proteoforms without mass shifts. (a) Distributions of the P1 residues for the C-termini of N-terminal and internal proteoforms. (b) Distributions of the P1 residues for the N-termini of C-terminal and internal proteoforms. (c) Distributions of the P1' residues for the C-termini of N-terminal and internal proteoforms. (d) Distributions of the P1' residues for the N-termini of C-terminal and internal proteoforms.

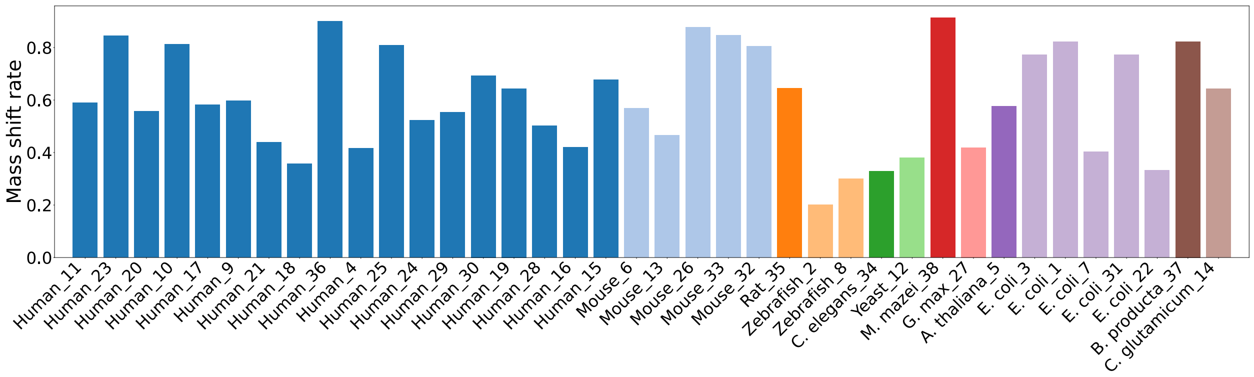


**Fig. S5**: Percentages of proteoform identifications with mass shifts across the 38 projects.

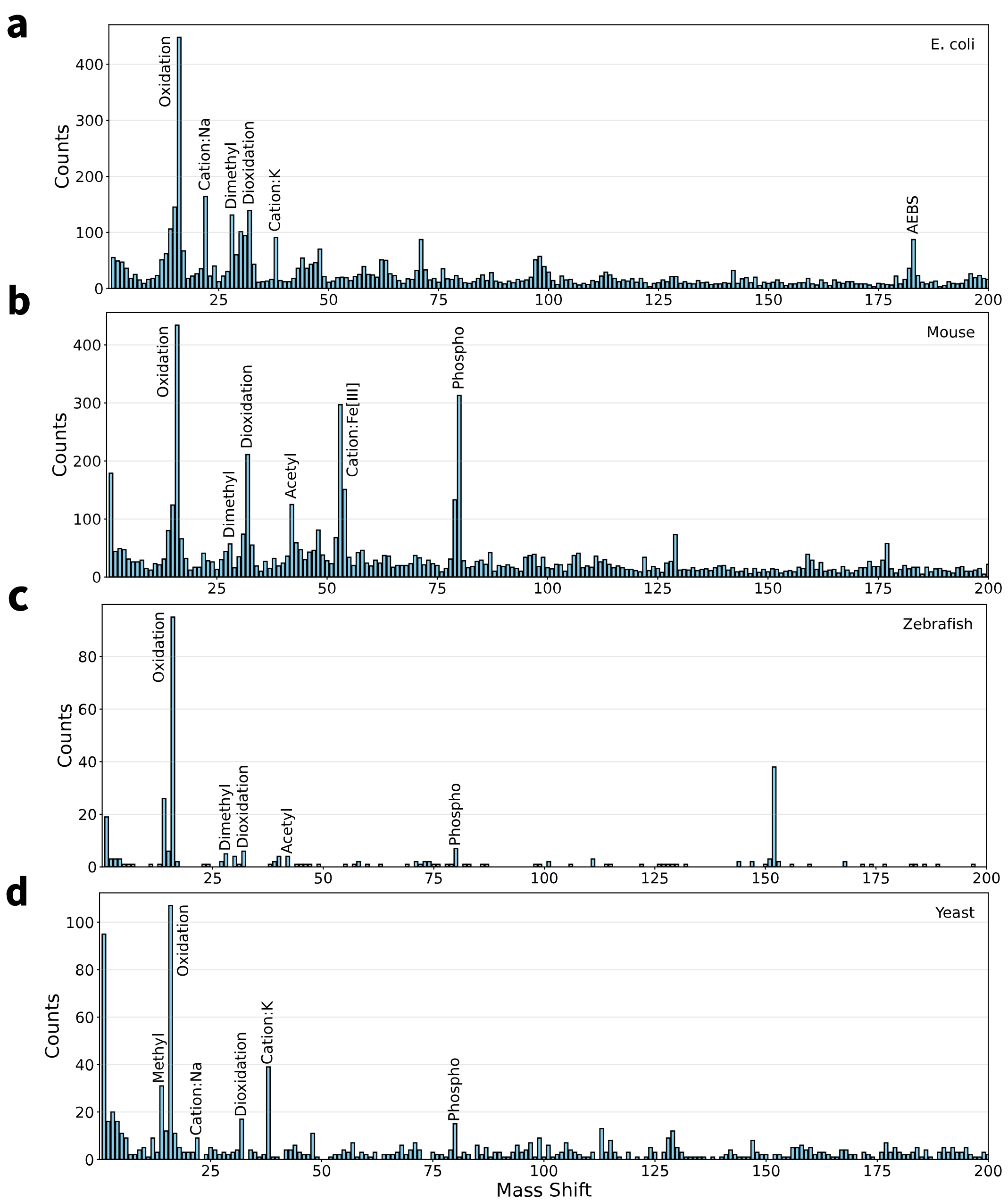


**Fig. S6:** Distributions of observed mass shifts in proteoform identifications in the range of [0-200] Da with a bin size of 1 Da for (a) *E. coli*, (b) mouse, (c) zebrafish, and (d) yeast.

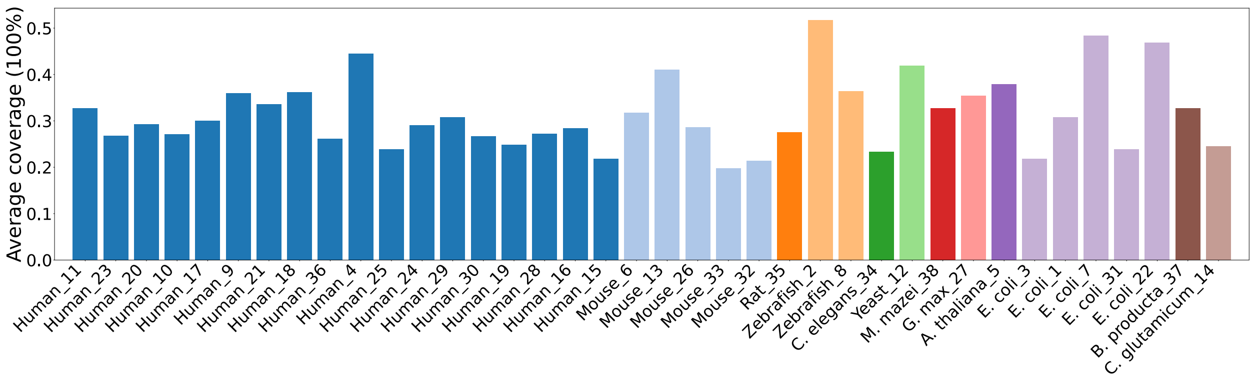


**Fig. S7**: Average sequence coverage of the proteoforms identified across the 38 projects.

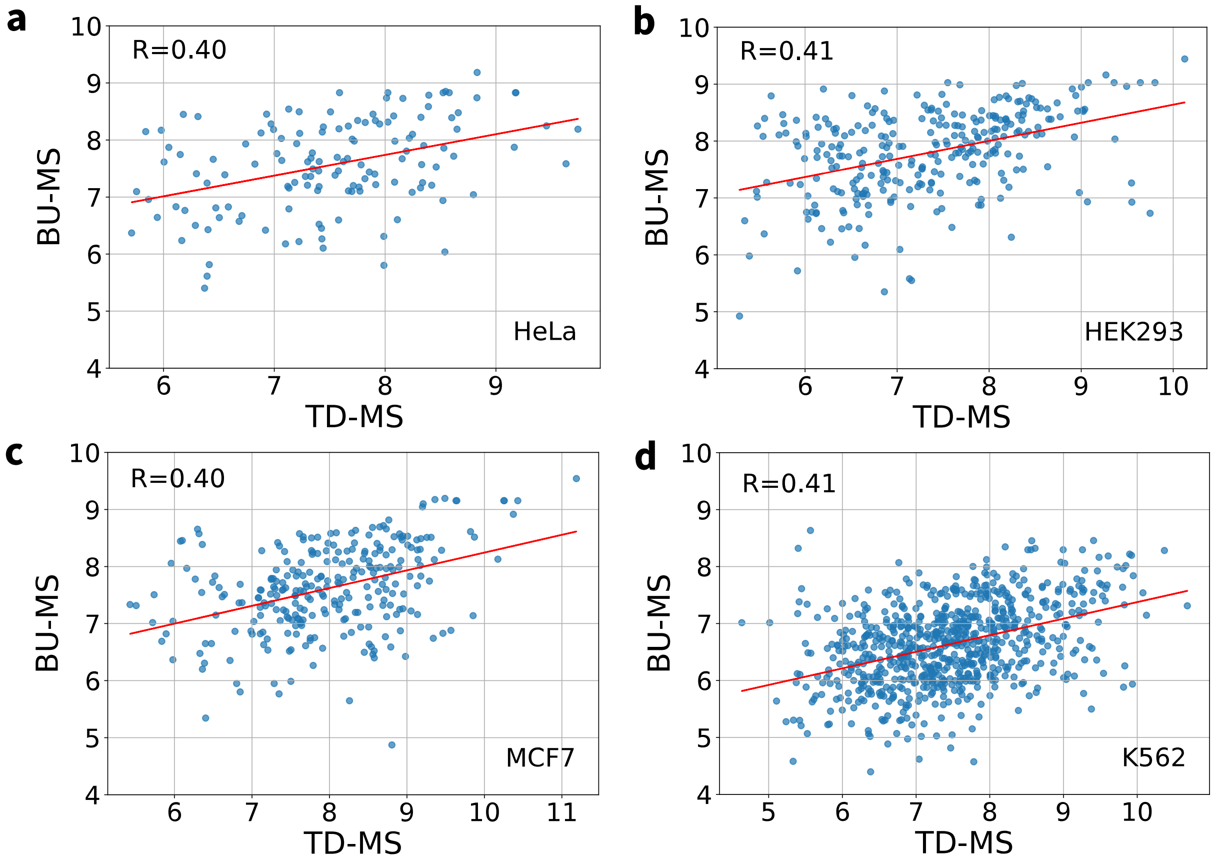


**Fig. S8:** Pearson correlations between protein abundances reported by TD-MS and BU-MS. (a) Hela (b) HEK293 (c) MCF-7 (d) K562. The red line indicates the fitted linear regression.

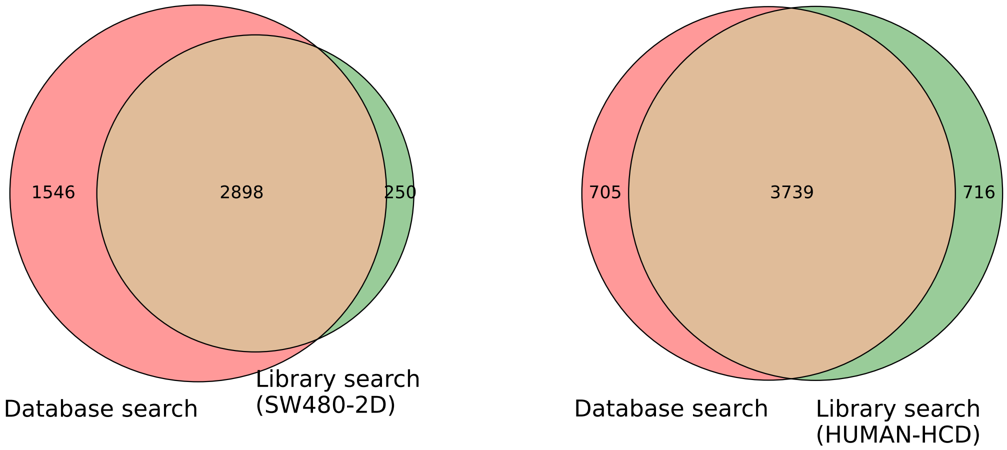


**Fig. S9:** Comparison of proteoform identifications from the second replicate of the SW480-2D dataset using database and spectral library searches. Proteoform identifications obtained by database search are compared with those obtained by spectral library search using (a) the SW480-2D library and (b) the HUMAN-HCD library.

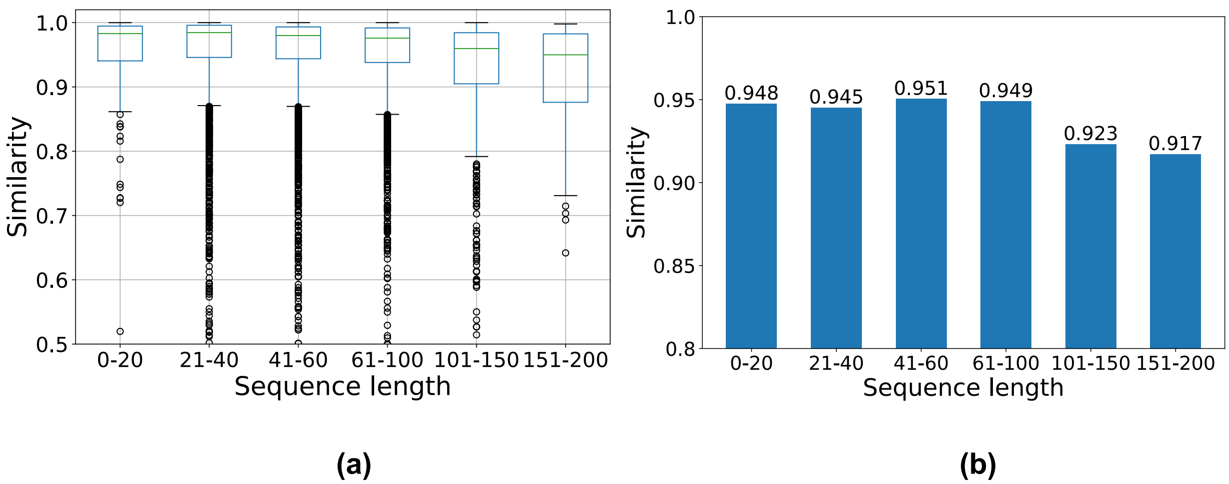


**Fig. S10:** Cosine similarity of 10,000 spectral pairs derived from identical proteoforms and charge states. (a) Distributions of cosine similarity for different proteoform lengths. (b) Average cosine similarity for different proteoform lengths. The average similarity of the 10,000 spectral pairs is 0.946.

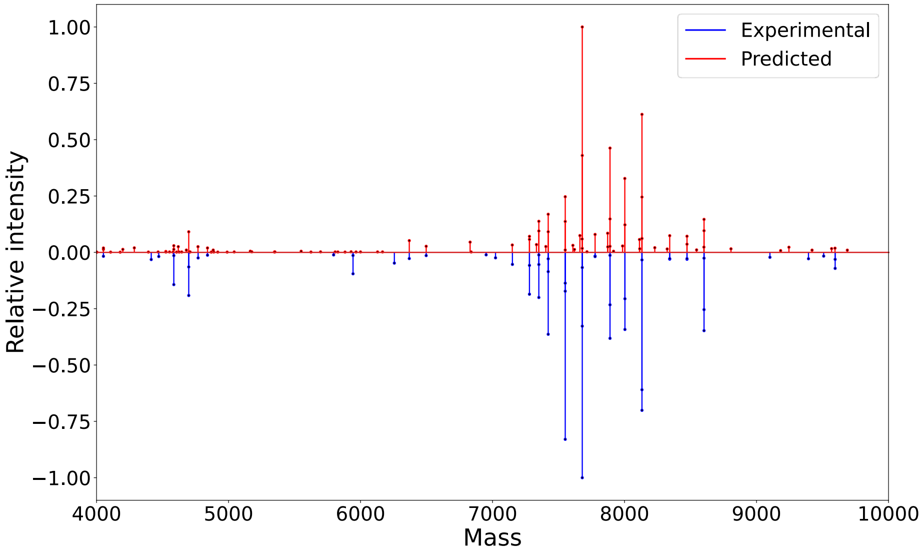


**Fig. S11:** An experimental deconvoluted spectrum and its corresponding predicted deconvoluted spectrum with a cosine similarity score of 0.938. Each peak in the deconvoluted spectrum represents the monoisotopic mass and the summed intensity of its corresponding isotopic envelope. The spectrum was generated from a proteoform of protein sp|P14174|MIF_HUMAN comprising 114 amino aicds (positions 2-115), with a monoisotopic mass of 12336.14 Da and a charge state of 8. The proteoform sequence is :PMFIVNTNVPRASVPDGFLSELTQQLAQATGKP PQYIAVHVVPDQLMAFGGSSEPCALCSLHSIGKIGGAQNRSYSKLLCGLLAERLRISPDRVYINY YDMNAANVGWNNSTFA.

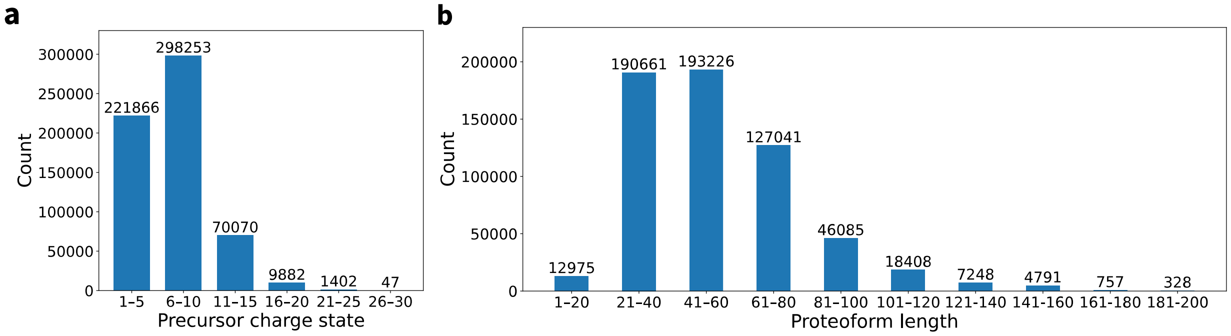


**Fig. S12:** The distributions of (a) precursor charge states and (b) proteoform length in the TRAIN-ALL data.

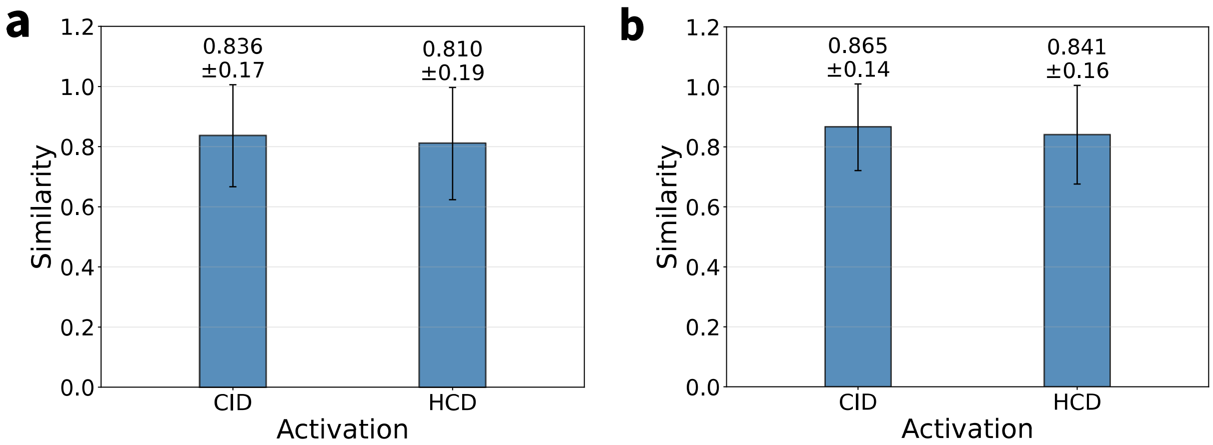


**Fig. S13:** Prediction accuracy of TD-Pred for HCD and CID spectra in the VAL_ALL dataset using (a) the backbone representation and (b) the simplified reprsentation.
